# Ultrafast and accurate 16S microbial community analysis using Kraken 2

**DOI:** 10.1101/2020.03.27.012047

**Authors:** Jennifer Lu, Steven L Salzberg

## Abstract

For decades, 16S ribosomal RNA sequencing has been the primary means for identifying the bacterial species present in a sample with unknown composition. One of the most widely-used tools for this purpose today is the QIIME (Quantitative Insights Into Microbial Ecology) package. Recent results have shown that the newest release, QIIME 2, has higher accuracy than QIIME, MAPseq, and mothur when classifying bacterial genera from simulated human gut, ocean, and soil metagenomes, although QIIME 2 also proved to be the most computationally expensive method. Kraken, first released in 2014, has been shown to provide exceptionally fast and accurate classification for shotgun metagenomics sequencing projects. Bracken, released in 2016, then provided users with the ability to accurately estimate species or genus abundances using Kraken classification results. Kraken 2, which matches the accuracy and speed of Kraken 1, now supports 16S rRNA databases, allowing for direct comparisons to QIIME and similar systems. Here we show that, using the same simulated 16S rRNA metagenomic data as previous studies, Kraken 2 and Bracken are up to 300 times faster and also more accurate at 16S profiling than QIIME 2.

## Background

Since the 1970s, sequencing of the 16S ribosomal RNA gene has been used for analyzing and identifying bacterial communities [1, 2]. This technology targets the 16S rRNA gene, which has regions that are both highly conserved and highly variable (hypervariable) among bacterial species. The highly conserved regions allow for the design of “universal” PCR primers to target and amplify the 16S sequence, while the hypervariable regions allow for discrimination among different bacterial clades. These properties allow 16S sequencing experiments to capture nearly all of the bacteria in a microbial community, which can then be compared to large 16S databases to determine their identities.

Researchers have utilized 16S rRNA sequencing for a very broad range of environmental and clinical studies. For example, the Earth Microbiome Project [3] and other environmental studies have used 16S sequencing to reveal the bacterial diversity of soil [4, 5], beach sand [6], and ocean environments [7]; while other studies targeted the microbiome of plants [8, ?, 9]. In the clinic, 16S rRNA has been used for diagnostic purposes to identify infectious bacterial species [10, 11, 12] and to characterize the role of bacterial diversity in diseases such as diabetes [13], Alzheimer’s disease [14], cancer [15], and autism [16]. The international Human Microbiome Project has used 16S data to characterize the bacterial community present in the human gut, feces, skin, and other areas of the body [17, 18, 19].

### 16S Classification

Analysis of the bacterial community from a 16S rRNA sequencing experiment involves comparing the reads to reference database. The tool most widely used for 16S classification today is the Quantitative Insights into Microbial Ecology (QIIME) software package [20], which compares sequencing reads against a 16S reference database. The three standard 16S databases, each of which has somewhat different content, are Greengenes [21], SILVA [22], and RDP [23].

First released in 2011, QIIME 1 [20] provided 4 classification algorithms for 16S rRNA, respectively based on the RDP classifier [24], BLAST [25], UCLUST [26], and SortMeRNA [27]. In 2018, QIIME 2’s q2-feature-classifier was released [28], adding 3 new classification algorithms based on scikit-learn’s naïve Bayes algorithm [29], VSEARCH [30], and BLAST+ [31]. By default, QIIME 1 uses the UCLUST algorithm for classification while QIIME 2 suggests usage of the naïve Bayes algorithm.

In 2018, Almeida et al. performed benchmark tests comparing QIIME 2 to its predecessor, QIIME 1, and to two additional 16S classification tools, MAPseq [32] and mothur [33]. Almeida et al. evaluated the performance of each tool by classifying 16S rRNA reads that were simulated from bacteria known to be present in human gut, soil, and ocean microbiomes. That study concluded that QIIME 2 provides the best accuracy on the basis of recall and F-score. However, they also noted that QIIME 2 was the most computationally expensive, requiring substantially more CPU time and more memory than other tools.

### Kraken, Kraken 2, and Bracken

The Kraken program uses an alignment-free algorithm that, when first released in 2014, was hundreds of times faster than any previously described program for shotgun metagenomics sequence analysis, with accuracy comparable to BLAST and superior to other tools [34]. Using a single thread, Kraken can classify metagenomics sequence data at a rate of >1 million reads per minute.

In 2016, Bracken was released as an extension to Kraken to estimate species abundance from Kraken’s output [35]. As originally designed, Kraken attempts to classify each read as specifically as possible, allowing reads to be classified at any taxonomic level depending on how many genomes share the same sequence. For example, a read that has identical matches to two species will be classified at the genus level. Bracken adds the capability of abundance estimation to Kraken; i.e., using Kraken’s read counts and prior knowledge of the database sequences, it estimates read counts for all species, genera, or higher-level taxa in a sample. For example, when Bracken is asked to estimate species counts, it will re-distribute all reads that Kraken assigns at the genus level (or higher) down to the species level.

Kraken 2, released in 2018, implemented significant changes to the database structure and classification steps to make databases smaller and classifications faster [38]. Because it uses the same classification algorithm, Kraken 2 has nearly the same precision and sensitivity as Kraken 1. However, Kraken 2 now also provides direct support for 16S classification with any of the three standard 16S databases: Greengenes, SILVA, and RDP. This new feature allowed us to compare Kraken 2 to the current state-of-the-art programs for 16S classification, as described below.

### Kraken 2 versus QIIME 2

In 2016, Lindgreen et al. evaluated 14 metagenomics classifiers, including Kraken 1 and QIIME 1 (UCLUST) [36]. That study showed that Kraken achieved the lowest false positive rate, 0%, while QIIME had a false positive rate of 0.28%. Kraken also had higher sensitivity than QIIME, correctly labeling 70% of the reads while QIIME was correct on 60%. Finally, Kraken obtained a Pearson correlation between the known and predicted abundances of phyla and genera of 0.99, versus 0.78 for QIIME. However, that study used different databases and different input data (reads produced by metagenomic shotgun sequencing) to evaluate these tools. For Kraken 1, Lindgreen et al. measured its performance on all input sequences from a shotgun metagenomics experiment, using a database containing all complete bacteria and archaeal genomes from RefSeq, while for QIIME 1, they analyzed its performance only on 16S rRNA sequences against the 16S Greengenes database.

Because QIIME is designed for 16S sequencing projects and Kraken has previously been used primarily for metagenomics shotgun sequencing projects, the tools have not been directly compared. Here, we compare QIIME 2 and Kraken 2 using the 16S rRNA reads generated in the Almeida et al. benchmark study [37], using both the Greengenes and SILVA 16S databases. We also show results for Kraken on the RDP database, which is not compatible with QIIME 2. Because we only tested the most recent version of each tool, we will henceforth refer to QIIME 2 as QIIME and Kraken 2 as Kraken.

## Results

Prior to classification, Kraken requires users to first build a specialized database consisting of three files: taxo.k2d, opts.k2d, and hash.k2d. The user also can choose the value *k* that determines the length of the sequences that Kraken uses for its index; every sequence (or k-mer) of length k is associated with the species in which it occurs. *K*-mers that occur in two or more species are associated with the lowest common ancestor of those species. The database files contain the taxonomy and k-mer information for the specified database. Following generation of these files, Bracken requires users to generate a k-mer distribution file. Kraken and Bracken additionally allow the use of multiple threads to accelerate database construction. We tested building all files for the 16S Greengenes 13_8, SILVA 132, and RDP 11.5 databases using 1, 4, 8, and 16 threads. **Table 1** summarizes the contents of each of these databases.

**Table 1.**
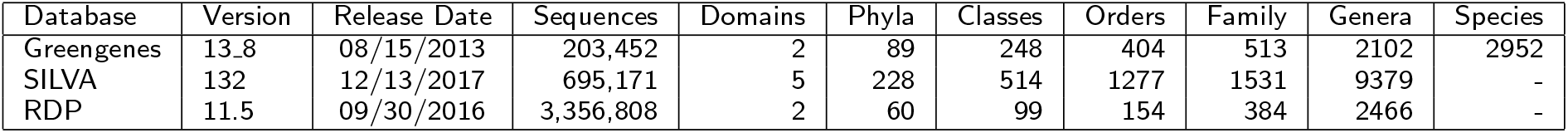
16S Databases used for the metagenomics classifiers in this study. For each of the most recently released versions of three 16S databases, this table describes the total number of sequences and the number of “traditional” nodes represented in their respective taxonomies. The Greengenes numbers refer to the 99% OTU database, and the SILVA values reflect the Ref NR 99 database.

For QIIME, users generate the database (called a “classifier”) by first converting sequence and taxonomy files into QIIME compatible .qza files. QIIME classifier generation is single-threaded. We built QIIME naïve-bayes classifiers for Greengenes 13_8 and SILVA 132.

**Figure 1A** compares the combined database building time for Kraken/Bracken against the classifier generation time of QIIME. Kraken was at least 9x faster than QIIME for database creation; e.g., it took 9 min to build the Greengenes database index, while QIIME required 78 minutes for the same database. For the SILVA database, Kraken required only 34 minutes while QIIME required more than 58 hours to build the same database. **Supplemental File 1** lists all command lines used for building the databases.

**Figure 1.**
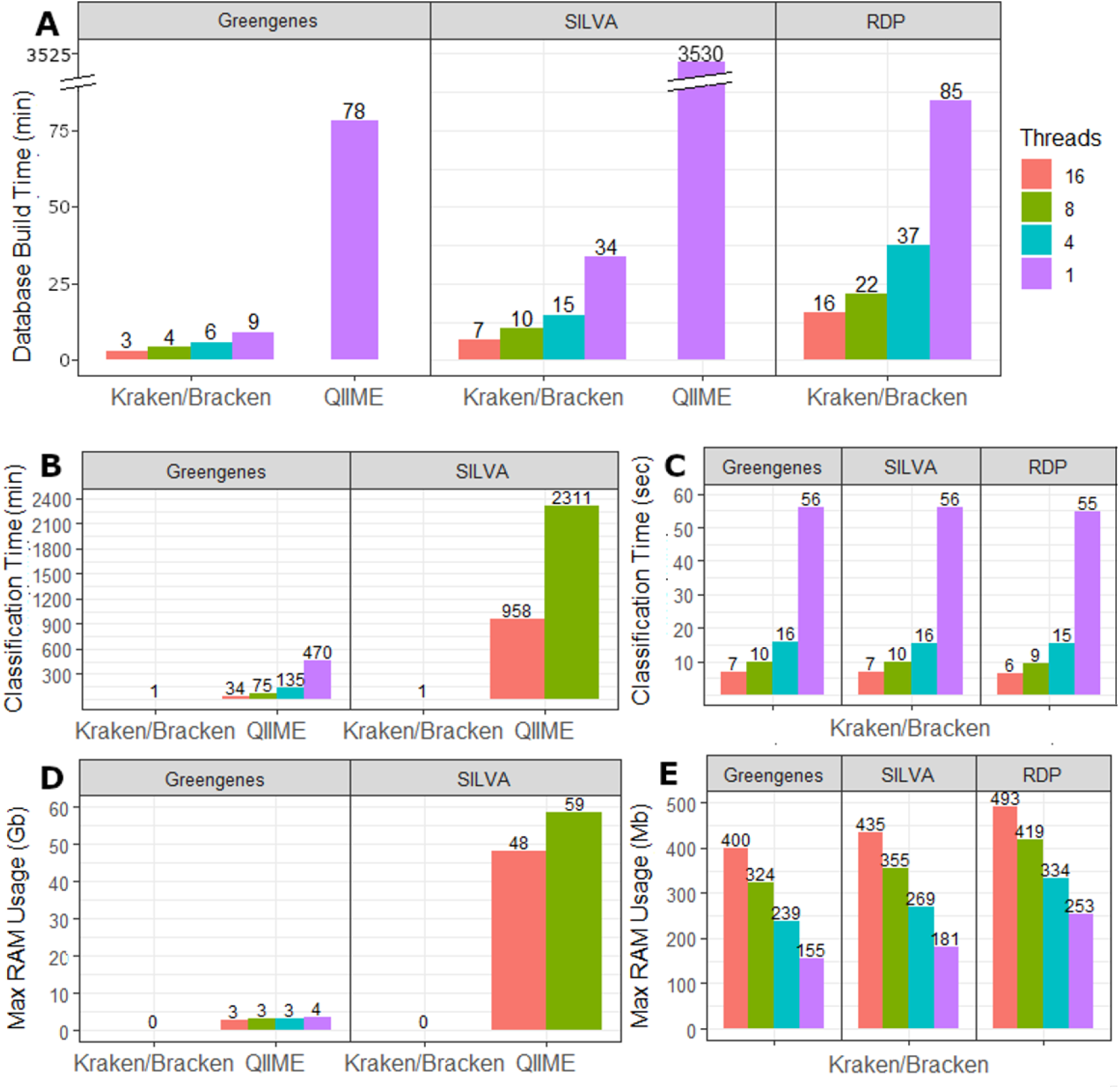
Build and Classification Statistics. **A)** Required time to build each database for Kraken/Bracken and QIIME. Kraken and Bracken allow for multi-threading while QIIME is single-threaded. **B)** Average classification runtime in minutes for each database. Kraken/Bracken combined runtime is reported for only 1 thread as all runtimes are < 1 min and bars are too small to be visible at this scale. QIIME was only run using 16 and 8 threads for SILVA. **C)** Classification runtime for Kraken and Bracken in seconds for all multi-threading options. **D)** Computational memory usage (RAM) for QIIME and Kraken/Bracken, shown in gigabytes (Gb). Kraken/Bracken RAM requirements reported only for 1 thread as Kraken and Bracken require < 0.5Gb of RAM regardless of thread count. **E)** Computational memory usage (RAM) for Kraken/Bracken shown in megabytes (Mb).

To compare the accuracy of Kraken, Bracken, and QIIME, we classified 12 samples generated by Almeida et. al. [37]. These 12 samples, each containing just under 200,000 reads, represent 3 different metagenomes (human, ocean, and soil) and 4 different 16S primers (V12, V34, V4, and V45). The number of reads in each sample is shown in **Table 2**.

**Table 2.** Sample Read Counts. The read counts in each metagenome-primer sample. Each sample was generated as described in the Supplementary Methods.

| Read Counts | V12 | V34 | V4 | V45 | Total |
| --- | --- | --- | --- | --- | --- |
| Human microbiome | 186,689 | 189,972 | 193,787 | 192,319 | 762,767 |
| Soil microbiome | 196,254 | 193,564 | 196,226 | 194,325 | 780,369 |
| Ocean microbiome | 193,867 | 193,962 | 196,198 | 195,135 | 779,162 |

QIIME classifiers require one single file containing all de-multiplexed reads. Therefore, we provided QIIME with one file per metagenome, each containing reads from all 4 primer sets. However, Kraken and Bracken classify samples one at a time, requiring each of the 12 samples to be processed individually.

Kraken and QIIME provide multi-threading options to speed up classification. We therefore tested Kraken and the QIIME Greengenes classifier using 1, 4, 8, and 16 threads. The QIIME SILVA classifier with 8 threads required approximately 1.5 days of run time, and for this reason we only tested it using 16 and 8 threads and did not evaluate the QIIME 2 SILVA classifier using 1 or 4 threads.

**Figure 1B** compares the average time in minutes required by QIIME vs. Kraken/Bracken to classify a single metagenome using the 16S Greengenes and SILVA databases. Due to the very large difference in run time between tools, this figure compares the multi-threaded options of QIIME against the single-threaded classification time of Kraken/Bracken. **Figure 1C** reports the classification times of Kraken/Bracken in seconds.

Another important consideration for software selection is the computational memory resources required. We evaluated this by measuring the RAM in gigabytes (GB) required for both classifiers. **Figure 1D** compares the RAM required for the single-threaded runs of Kraken/Bracken against the multi-threaded runs using QIIME. Notably, all Kraken/Bracken runs used less than 0.5 GB of RAM, which appears in the figure as zero GB. To provide more detail on RAM usage, **Figure 1E** reports the RAM required by Kraken/Bracken in megabytes (MB) for all multi-threading options.

The resulting counts per genus for each of the human, ocean, and soil samples are listed in **Supplemental Tables 1, 2, and 3**, respectively. **Figure 2** compares the true distribution of genera in each metagenomics sample against the genus-level counts reported by Kraken 2, Bracken, and QIIME 2. For clarity, this figure shows the combined read counts across the V12, V34, V4, and V45 samples for each metagenome.

**Figure 2.**
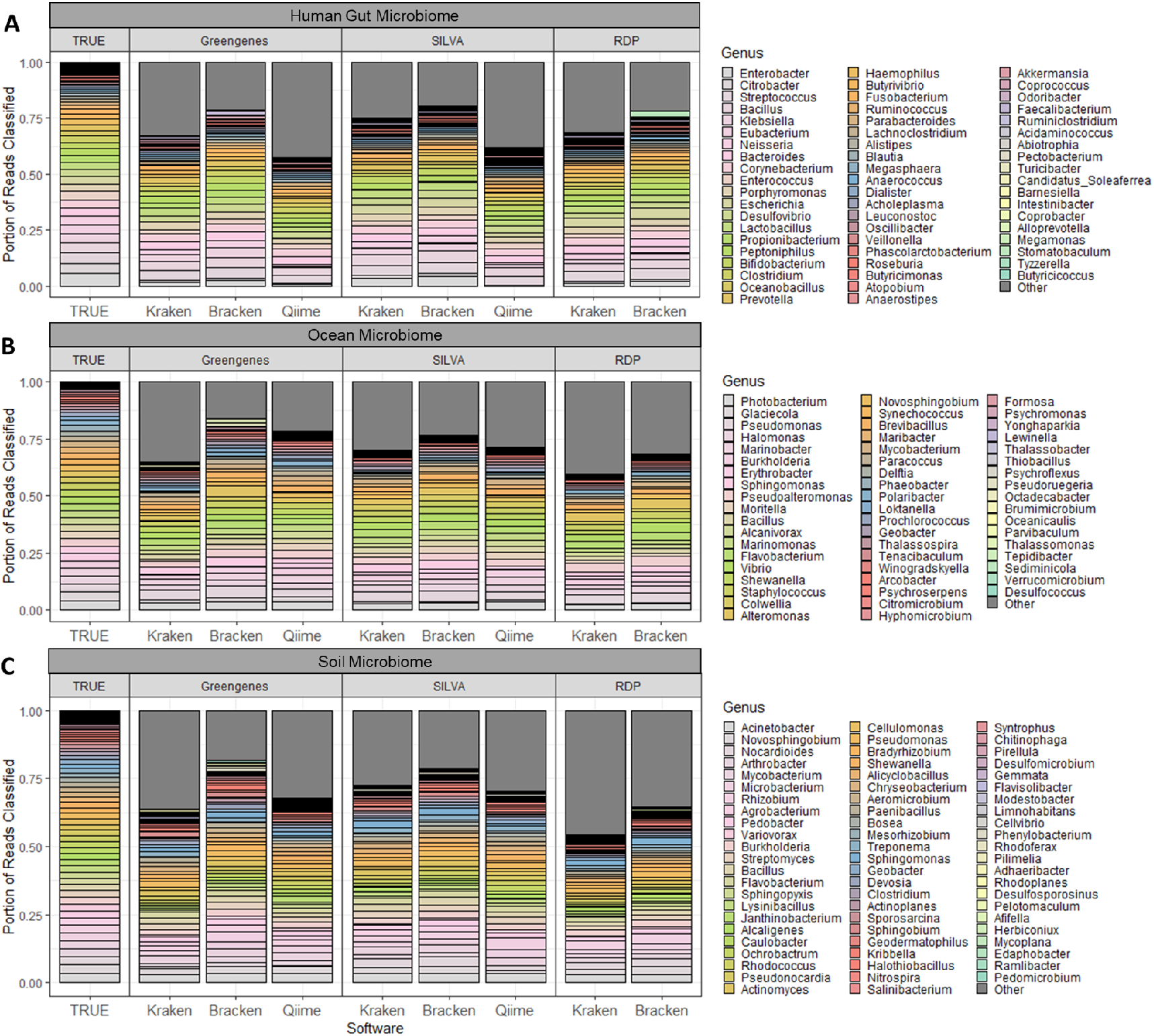
Genera Distribution for Simulated Microbiota. This plot compares the true genus abundances against those abundances estimated by Kraken, Bracken, and QIIME, for each of the three simulated microbiome samples (**A** = human gut microbiome, **B** = ocean microbiome, **C** = soil microbiome). Only the correct genera are represented by different bars while read assignments to any incorrect taxon is included in “Other”.

We used two different metrics to evaluate the genus distribution accuracy: Mean Absolute Percentage Error (*MAPE*) and Bray-Curtis dissimilarity (*BC*). Both error rates measure how different the predicted sample distribution is from the true genera counts. See the **Methods** section for details on how each error rate is calculated. Given these two metrics, we evaluate accuracy as 1 – *MAPE* and 1 – *BC*. **Figure 3A** compares the accuracy of each tool when calculating the correct combined read counts at the genus level for each metagenome. For further insight into how the choice of 16S primer affects genus distribution accuracy, we evaluated the average *MAPE* and average *BC* across all 3 metagenome samples for each program/database. **Figure 3B** uses these averages to compare the accuracy between 16S primers. **Supplemental Table 4** lists all *MAPE* and *BC* values for each combination of software/database/primer/metagenome.

**Figure 3.**
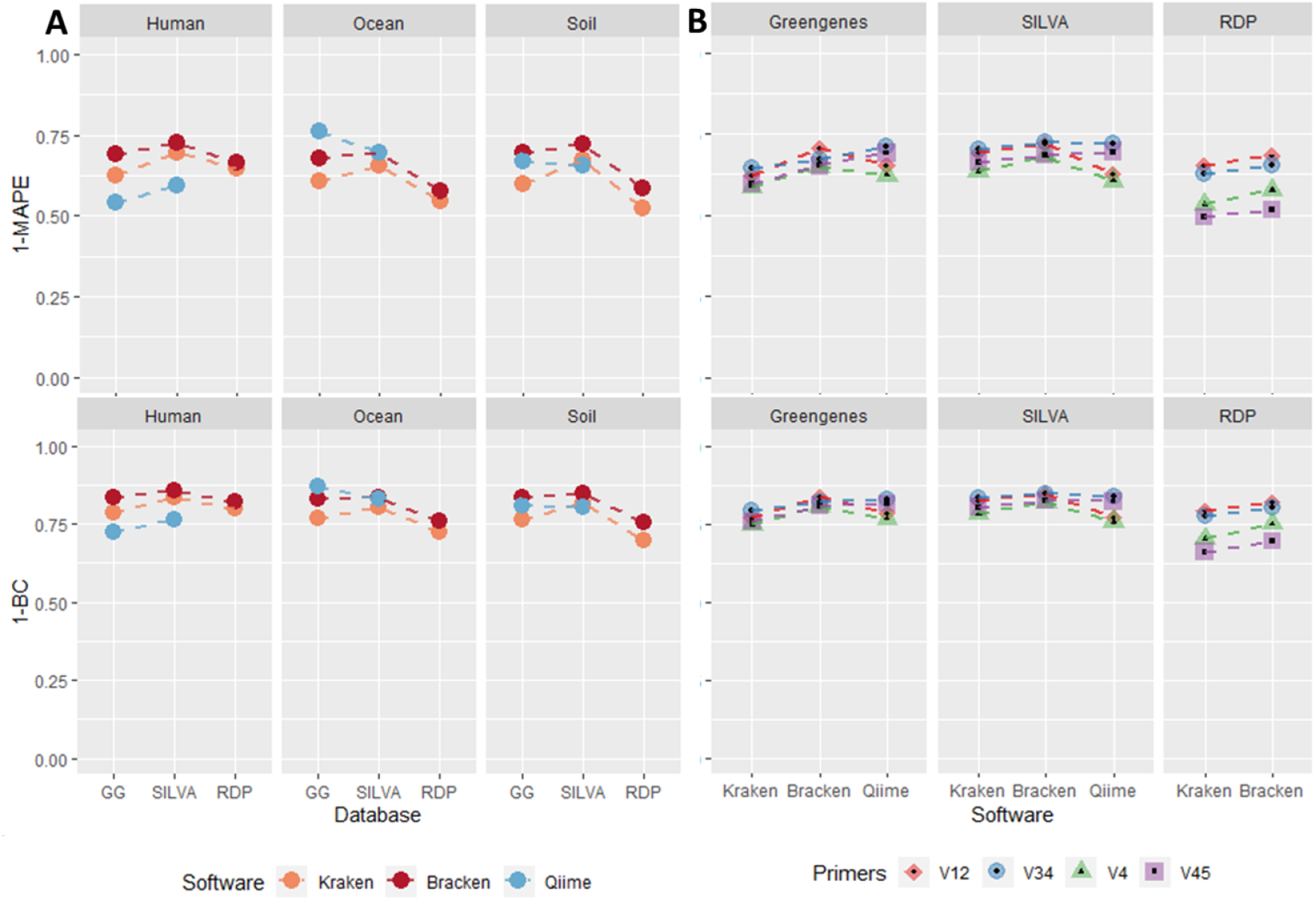
MAPE and Bray-Curtis Dissimilarity. This plot evaluates classification accuracy by using the inverse of two error metrics: Mean Absolute Proportion Error (*MAPE*) and Bray-Curtis Dissimilarity (*BC*). **A** compares the accuracy of Kraken, Bracken, and Qiime when predicting the genus read counts across all samples for given metagenome/database. **B** compares the accuracy between the individual primers averaged across all 3 metagenomes for a given software/database. The top plots calculate accuracy as 1 – *MAPE* while the bottom plots evaluate 1 – BC.

While all tools tested provide general read counts per genus, Kraken is the only tool that directly assigns each read with a taxonomic label. Using this information, we can calculate Kraken’s accuracy when classifying reads at major taxonomic levels in terms of sensitivity and precision. We measure precision by positive predictive value (PPV, see Supplemental Methods for more details). **Figure 4** displays Kraken’s average sensitivity and PPV for each database used (**Figure 4A**) and for each 16S primer used in generating the samples (**Figure 4B**).

**Figure 4.**
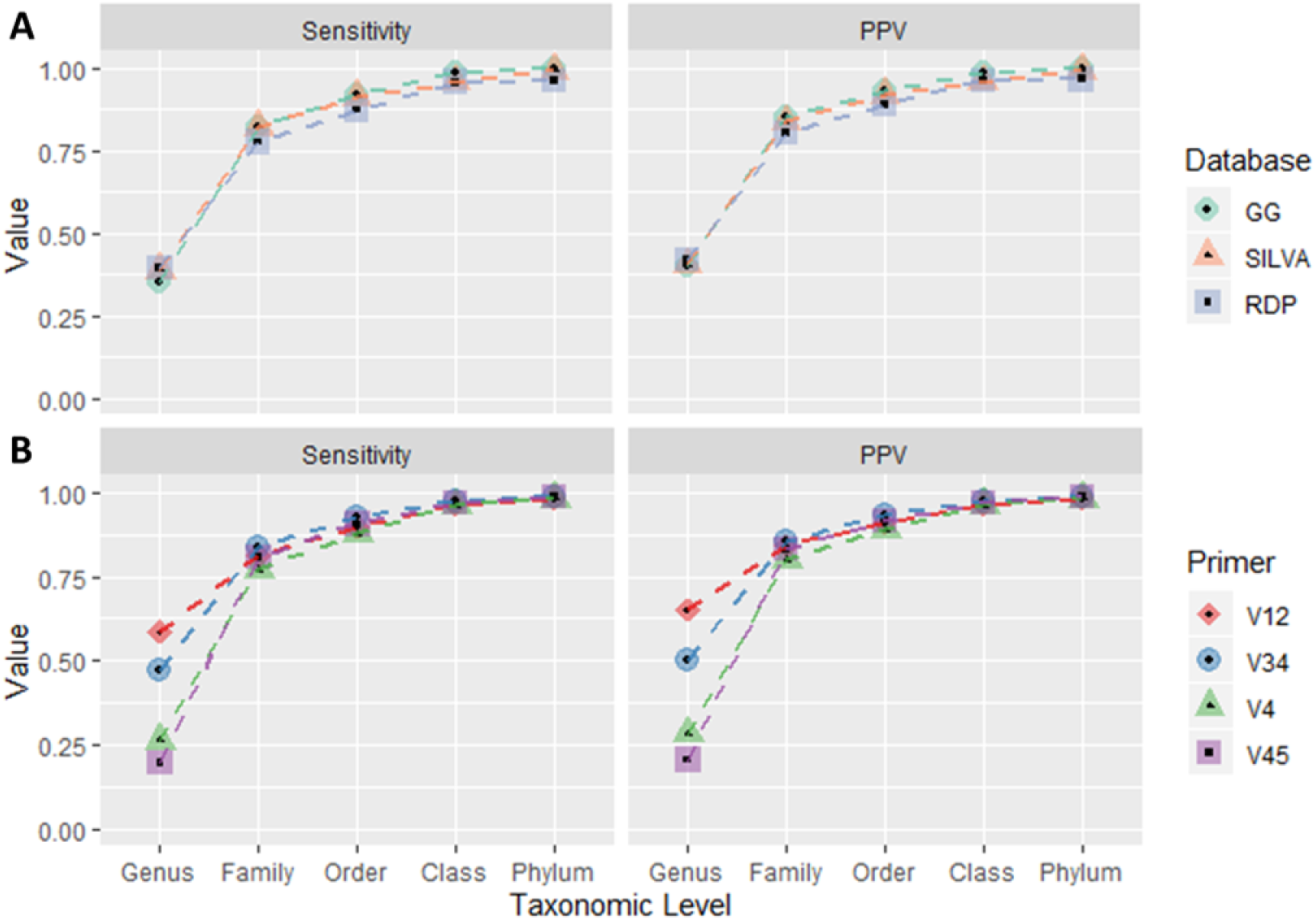
Kraken Per-Read Accuracy. As Kraken is the only tool tested that provides per-read taxonomy assignments, we evaluate the sensitivity and precision (*PPV*) of Kraken 2’s taxonomy assignments at each major taxonomic level

## Discussion

In this study, we evaluated three systems for classification and abundance estimation of 16S sequencing data sets: Kraken 2, Bracken, and QIIME 2. For Kraken and Bracken, we used three 16S databases: Greengenes, SILVA, and RDP; while for QIIME, we only evaluated Greengenes and SILVA. We then used these tools/databases to classify 12 samples generated by Almeida et. al [37], which represent 3 simulated metagenomes (human gut, ocean, and soil) and 4 different 16S primers (V12, V34, V4, and V45). In total, we collected 36 different results using Kraken/Bracken and 24 different results using QIIME.

### Database Building Time

For all systems compared here, database build time is a function of the number of sequences in the database. Because 16S Greengenes is the smallest database (with 200,000 sequences) and 16S RDP is the largest (with 3.4 million sequences), generation of database files is fastest with Greengenes and slowest with RDP.

When comparing single-threaded Kraken/Bracken against QIIME, Kraken and Bracken combined require far less build time. For the smallest 16S database, Greengenes, QIIME required more than an hour to generate the naïve Bayes classifier (**Figure 1A**). By comparison, single-threaded Kraken and Bracken combined required less than 10 minutes to create the database files. For 16S SILVA, with nearly 700,000 sequences, QIIME 2 required more than 58 hours for classifier generation while the single-threaded Kraken/Bracken required only ~30 minutes. We additionally note that the largest 16S database, RDP, required a little more than an hour for single-threaded Kraken 2 and Bracken to create the database files. As mentioned above, the RDP database is incompatible with QIIME 2. The multi-threaded nature of Kraken 2 and Bracken further accelerate the database building process, with 4 threads halving the required build time (**Figure 1A**).

### Classification Time/Memory Requirements

As observed by Almeida et. al. [37], QIIME 2 requires more computational resources than other methods during classification. With the use of 16 CPU threads, QIIME required 35 minutes on average to classify the human, ocean, and soil metagenomic samples using the Greengenes database (**Figure 1B**). The QIIME’s SILVA classifier required 16 hours on average. By comparison, single-threaded Kraken 2 and Bracken required on average 1 minute per metagenomic sample. This runtime decreases from 1 minute to 15, 10, and 6 seconds for 4, 8, and 16 threads respectively (**Figure 1C**). The runtime of Kraken 2 and Bracken was nearly the same for all three databases. Thus Kraken or Braken is at least 350 times faster (6 seconds vs. 35 minutes) than QIIME 2 when run with 16 parallel threads.

The amount of computer memory (RAM) required by each system also varied widely (**Figure 1D**). For all three databases, single-threaded Kraken required < 260 MB of RAM. However, the single-threaded QIIME Greengenes classifier required 3.6 GB of RAM. Increasing the number of threads for Kraken also increases the total RAM used, with 16 threads using 400-500 MB of RAM for each of the Kraken databases (**Figure 1E**). However, for QIIME, increasing the number of threads decreased the total RAM: the QIIME Greengenes classifier with 16 threads used ~2.7 GB, and the QIIME SILVA classifier with 16 threads used 48 GB of RAM (**Figure 1D**).

### Accuracy of abundance estimation

Finally, we compared the accuracy of all three tools based on their ability to recreate the true genus distribution of the simulated samples (**Figure 2**). We quantified the accuracy of these distributions using both MAPE and Bray-Curtis dissimilarity (**Supplemental Table 4**).

In all cases, Bracken performed better than Kraken 2, which was expected because Kraken is a classification tool, not an abundance estimation system. Kraken classifies reads at any level in the taxonomy, which means that some reads might be assigned to a higher level genus; e.g., any read that has equally good matches to two genera will be assigned to the family containing them. For the simulated datasets in this study, Kraken assigned from 7-30% of the reads to levels above genus. These reads are not incorrectly classified, but the result is that Kraken underestimates the abundances of their genera. By contrast, Bracken is designed to use Kraken’s classification data to estimate all read counts at the genus level, thereby improving on Kraken’s genus-level distribution.

On average, Bracken performed the best, having the lowest average error rates across all three 16S databases (**Supplemental Table 4**). Bracken also had the lowest error rate for 8/9 combinations of samples and databases. The only sample where QIIME 2 had a lower error rate than Bracken was in the classification of the ocean samples against the 16S Greengenes database (**Figure 3A**). However, QIIME 2 had the highest error rate when classifying the human sample against Greengenes or SILVA, regardless of whether measured by MAPE or Bray-Curtis dissimilarity.

In analyzing the trends across the databases using both MAPE and Bray-Curtis, Bracken performed the best using the 16S SILVA database and performed the worst using the 16S RDP database. 16S RDP yielded on average 0.391 MAPE and 0.221 BC Index while 16S SILVA only yielded a 0.286 MAPE and a 0.153 BC Index. 16S Greengenes with Bracken had an average of 0.313 MAPE and a 0.165 BC Index. Although QIIME 2 was not tested on 16S RDP, QIIME 2 yielded the same trends when comparing 16S Greengenes and SILVA, with 16S SILVA outperforming 16S Greengenes in almost all cases.

In addition to evaluating the different tools, we also evaluated the accuracy of each of the primer sets (V12, V34, V4, and V45) that were used by Almeida et. al. [37]. **Figure 3B** shows the average accuracy of each primer set across all 3 metagenomes for a given software/database pairing. For both Greengenes and SILVA, the samples generated using V34 and V12 performed slightly better. However, for RDP, the difference in accuracy between primer samples is further magnified. When classifying with the RDP database, both Kraken and Bracken had significantly better results for the V12 and V34 samples (**Figure 3B**).

### Per-Read Classification Accuracy

Kraken is the only program of the three tested here that provide per-read assignments by default, allowing us to compute the read-level accuracy of its taxonomy assignments. Per-read accuracy is somewhat dependent on the reference database, but highly dependent on the 16S primer set (**Figure 4B**). In particular, Kraken had three times higher sensitivity (60%) and PPV (65%) when classifying reads generated using V12 primers versus those generated from V45 primers (20% and 21%).

As expected, sensitivity and precision increased with taxonomic level, with class and phylum sensitivity and precision exceeding 0.95 for all sample sets and all databases. **Supplemental Table 6** contains exact numbers for sensitivity and precision for each dataset and database.

### Taxonomy Inconsistencies

In our experiments, we discovered that the accuracy of 16S analysis is highly dependent on the choice of 16S database. The 170 distinct genera present in our human, ocean, and soil metagenomes were selected from the NCBI taxonomy, but none of the three 16S database taxonomies contains precisely the same genera. Each 16S database is independently curated from different reference sets, resulting in substantial differences among the taxonomies [38]. Among the 170 unique genera uses here, 22 are missing from Greengenes, 19 have different names or are mapped to multiple genera in RDP, and 16 have different names in Silva (see **Supplemental Table 5**). For example, Agrobacterium, Burkholderia, and Rhizobium are not unique genera in the 16S SILVA taxonomy, but are combined into a single “Allorhizobium-Neorhizbium-Pararhizobium-Rhizobium” genus. Escherichia and Shigella are also combined into the “Escherichia-Shigella” genus in 16S SILVA. The Clostridium sequences in 16S SILVA are split between 19 different genera, each with the prefix of “Clostridium sensu stricto” followed by a number 1-19.

## Conclusion

Although each of the 16S databases represents a large number of bacterial organisms, the accuracy of metagenomics classifiers varied substantially among them. In our experiments, 16S SILVA provided the lowest error rates and highest per-read accuracy regardless of the software used in classification. Across all databases, Kraken 2 and Bracken outperformed QIIME 2 in terms of computational requirements, runtime, and accuracy. Single-threaded Kraken/Bracken was nearly 8x faster than QIIME 2 at building the 16S Greengenes database and 100x faster at building a 16S SILVA database. Kraken and Bracken also allow for multi-threaded database building, which allows any 16S database to be built in less than 20 minutes. For classification, Kraken/Bracken used 20 times less RAM, performed 300 times faster, and achieved better genus-level resolution than QIIME 2.

## Methods

### Almeida Simulated Data

QIIME 2, Kraken 2, and Bracken were evaluated using the A500 synthetic microbiome samples generated by Almeida et al. [37] and available at ftp://ftp.ebi.ac.uk/pub/databases/metagenomics/taxon_benchmarking/. The A500 set contains 12 samples representing three different microbial environments: the human gut, ocean, and soil. For each of these environments, genomic sequences for their most abundant genera were extracted and randomly sampled. These human gut, ocean, and soil genomes then were sub-sampled four times to simulate 16S rRNA profiling using four different primer sets, generating 200,000, 250-bp paired-end reads per primer sequence. The sub-sampling introduced a 2% random mutation to each sequencing read. Almeida et al. then performed pre-processing and quality control to filter sequences with ambiguous base calls. With three microbial environments and four primer sets, Almeida et. al. thereby generated 12 sets of synthetic communities for testing. Information about the software and primers used in dataset generation is further described in the Methods section of Almeida et al.

### Software and Databases

The software packages tested are Kraken 2 (downloaded on 2020/03/05), Bracken v2.5 and QIIME 2 v2017.11. Kraken and Bracken database files were generated for Greengenes 13_8, SILVA 132, and RDP 11.5 database releases. QIIME 2 database files were generated for Greengenes 13_8 and SILVA 132.

### Error Rate Calculations

For evaluating the accuracy of Kraken 2, Bracken, and QIIME 2, we calculated two different error metrics which compare the true genera distributions against those reported by each program. The first error metric is a modified mean absolute proportion error (MAPE) which compares the difference between the true read counts (*T_g_*) for a given genus and the measured read counts (*A_g_*) for that same genus.

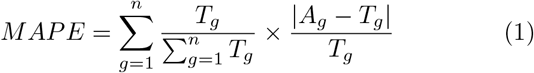

Each difference is calculated as a fraction of the true counts and then weighted by the fraction of the total sample. *n* is the total number of true genera in the sample.

The second metric, Bray-curtis dissimilarity [39], is a similar measurement of the dissimilarity between the true genera distribution and the measured genera distribution. The formula for Bray-curtis dissimilarity is:

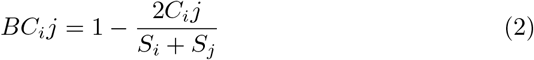

where *C_ij_* is the sum of lesser reads for genera in common and *S_i_* = *S_j_* is the total number of reads. In other words, for every true genus *g* in the sample, if *T_g_* < *A_g_, C_i_j* = *C_i_j* + *T_g_*. Otherwise if *T_g_* > *A_g_, C_i_j* = *C_i_j* + *A_g_*.

*MAPE* and *BC* values both fall between 0 and 1, where larger values indicate a greater difference between samples and smaller values indicate a greater similarity.

### Sensitivity and Precision (PPV) Calculations

As Kraken 2 provides taxonomic assignments for every read, we can use the true taxonomic tree of each read to calculate sensitivity and precision at all taxonomic levels. For this explanation, we describe our calculations of sensitivity and precision for the genus level. First, we calculate true positive (*TP*), vague positive (*VP*), false positive (*FP*), and false negative (*FN*) read counts. We define *TP* read counts as the number of reads correctly classified at the genus level. This includes reads that are classified as any species within the true genus. Vague positive (*VP*) reads account for the possibility that a read is classified as any ancestor of the true taxon. Therefore, *VP* reads include all *TP* reads and all reads assigned to ancestor taxa of the true genus. *FN* reads are all classified reads that are not *VP* reads. This thereby includes reads classified at any taxa not within the direct lineage of the true genera. Finally, we define *FN* as the number of unclassified reads. Notably, in all experiments, Kraken 2 did not label any read as unclassified (*FN* = 0).

From these values, we define sensitivity and precision (measured by positive-predictive-value, *PPV*) using the following two equations:

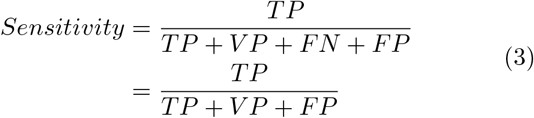

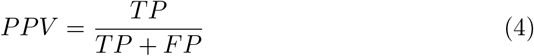

## Supporting information

Supplemental Information: Command Lines and Qiime Per Read Discussion

Supplemental Table 1: Human Microbiome Read Counts

Supplemental Table 2: Ocean Microbiome Read Counts

Supplemental Table 3: Soil Microbiome Read Counts

Supplemental Table 4: MAPE and Bray Curtis Errors

Supplemental Table 5: 16S Database Inconsistencies

Supplemental Table 6: Kraken Per Read Accuracy

## Abbreviations

*GG*: Greengenes
*TP*: true positive
*TN*: true negative
*FP*: false positive
*FN*: false negative
*MAPE*: Mean Absolute Percentage Error
*BC*: Bray-Curtis Dissimilarity
*PPV*: Positive Predictive Value

## Competing interests

The authors declare that they have no competing interests.

## Author’s contributions

JL conceived and designed the experiments, performed the experiments, analyzed the data, wrote the paper, prepared figures and/or tables, performed the computation work, and wrote the paper.

SLS conceived and designed the experiments, analyzed the data, and wrote the paper.

## Funding

This work was supported in part by NIH under grants R01-HG006677 and R35-GM130151 and by NSF under grant IOS-1744309.

## Supplemental Files

**SupplementalJnformation.pdf**: This file contains all command lines used for testing Kraken 2, Bracken, and QIIME 2 and with building each of the 16S databases. Additionally, this file contains a short discussion of the multi-threading behavior of QIIME 2.

**Supplemental_Table_1.xlsx**: Human Genus Read Counts – True v Kraken v Bracken v QIIME

**Supplemental_Table_2.xlsx**: Ocean Genus Read Counts – True v Kraken v Bracken v QIIME

**Supplemental_Table_3.xlsx**: Soil Genus Read Counts – True v Kraken v Bracken v QIIME

**Supplemental_Table_4.xlsx**: MAPE and Bray-Curtis Dissimilarities

**Supplemental_Table_5.xlsx**: 16S Database Inconsistencies

**Supplemental_Table_6.xlsx**: Kraken 2 Per Read Sensitivity and Precision

