## Supplemental Information: Command Lines and Qiime Per Read Discussion for "Ultrafast and accurate 16S microbial community analysis using Kraken 2"

QIIME 2 Results Affected by Thread Count **3**

#### Kraken 2 and Bracken Command Lines

##### 1. Database Download/Building

The following two command lines were executed for each of the 16S databases (greengenes, silva, rdp), using multi-threading options of 1, 4, 8, and 16 threads. As read lengths tested in this experiment are all 250bp, we used option `-l 250` for building the Bracken files.

```
kraken2-build --db ${DBNAME} --special ${DBTYPE} --threads ${THREADS}
bracken-build -d ${DBNAME} -t ${THREADS} -l 250 -k 35
```

##### 2. Classification/Abundance Estimation Steps

For each sample and each database combination, we executed the following two commands tested using 1, 4, 8, and 16 threads.

```
kraken2
    --db ${DBNAME}
    --threads ${THREADS}
    --report ${SAMPLE}.kreport2
    --paired ${SAMPLE}_R1.fa ${SAMPLE}_R2.fa > ${SAMPLE}.kraken2
bracken -d ${DBNAME} -r 250 -l G -i ${SAMPLE}.kreport2 -o ${SAMPLE}_g.bracken
```

#### QIIME 2 Command Lines

All QIIME 2 command lines were executed within a conda environment. Command lines are displayed using multiple lines for clarity.

##### 1. Import Sequence and Taxonomy Files

QIIME 2 compatible sequences for Greengenes 13_8 were downloaded from `ftp://greengenes.microbio.me/greengenes_release/gg_13_5/gg_13_8_otus.tar.gz`. The following command lines then were used to convert the files into QIIME 2 compatible files.

```
qiime tools import
    --type 'FeatureData[Sequence]'
    --input-path gg_13_8_otus_99.fasta
    --output-path gg_13_8_otus_16S.qza
qiime tools import --type 'FeatureData[Taxonomy]'
```

```
        --input-format HeaderlessTSVTaxonomyFormat
        --input-path 99_otu_taxonomy.txt
        --output-path gg_taxonomy.qza
```

QIIME 2 compatible sequences for SILVA 132 were downloaded from `https://www.arb-silva.de/download/archive/qiime/SILVA_132_release.zip`

```
    qiime tools import
        --type 'FeatureData[Sequence]'
        --input-path silva132_99.fna
        --output-path 99_otus_silva.qza
    qiime tools import
        --type 'FeatureData[Taxonomy]'
        --input-format HeaderlessTSVTaxonomyFormat
        --input-path 7_level_taxonomy.txt
        --output-path silva_taxonomy.qza
```

#### 2. Train Classifier

```
    qiime feature-classifier fit-classifier-naive-bayes
        --i-reference-reads gg_13_8_otus_16S.qza
        --i-reference-taxonomy gg_taxonomy.qza
        --o-classifier classifier_gg13_8.qza

    qiime feature-classifier fit-classifier-naive-bayes
        --i-reference-reads 99_otus_silva.qza
        --i-reference-taxonomy silva_taxonomy.qza
        --o-classifier classifier_silva132.qza
```

#### 3. Import Sample

QIIME 2 requires that all sample files first be converted into QIIME-compatible .qza files and that all sequences be dereplicated, generating a single sample file.

```
    qiime tools import --type ‘SampleData[Sequences]’
        --input-path combined_seqs_${SAMPLE}.fna
        --output-path  combined_seqs_${SAMPLE}.qza
    qiime vsearch dereplicate-sequences
        --i-sequences combined_seqs_${SAMPLE}.qza
        --o-dereplicated-table table-${SAMPLE}.qza
        --o-dereplicated-sequences rep_seqs_${SAMPLE}.qza
```

#### 4. Classification and Exporting Steps

The following command lines classifies the sample using the given trained classifier. This example uses the 16S Greengenes classifier. Additionally, this step allows for multi-threading. Therefore, we executed this command using 8 and 16 threads.

```
    qiime feature-classifier classify-sklearn
        --i-classifier classifier_gg13_8.qza
        -i-reads rep_seqs_${SAMPLE}.qza
        --o-classification ${SAMPLE}_classsified_gg.qza
        --verbose
        --p-n-jobs ${THREADS}
```

Following classification, the QIIME file must be exported using the following steps.

```
qiime tools export
    --input-path qiime_data/table-${SAMPLE}.qza
    --output-path ${SAMPLE}/
qiime tools export
    --input-path qiime_classified/${SAMPLE}_classified_silva.qza
    --output-path ${SAMPLE}/
```

Following these two steps, we changed the first line of taxonomy.tsv to "#OTUID taxonomy confidence" separated by tabs

```
biom add-metadata
    -i ${SAMPLE}/feature-table.biom
    -o ${SAMPLE}/${SAMPLE}-table-taxonomy.biom
    --observation-metadata-fp ${SAMPLE}/taxonomy.tsv
    --sc-separated taxonomy
biom convert
    -i ${SAMPLE}/${SAMPLE}-table-taxonomy.biom
    -o ${SAMPLE}/${SAMPLE}-table-taxonomy.tsv
    --to-tsv
    --header-key taxonomy
```

### QIIME 2 Results Affected by Thread Count

During our experiments, we discovered that changing the multi-threading options of QIIME 2's classification step affected the final results. Therefore, we tested the QIIME 2 16S Greengenes classifier using 1, 2, 4, 6, 8, 12, 16, 20, 24, 28, 30, and 32 threads and the human-gut microbiome sample. The results indicated that all results using 6-32 threads were identical, but different from the results using 1-4 threads. Additionally, we then were unable to replicate these results when executing the QIIME classifier commands at a different time.

Specifically, we calculated the MAPE error for the QIIME Greengenes classifier classifying the human gut microbiome samples. The MAPE calculated from the initial test using 8 threads was 0.459. However, when we ran the classifier consecutively for 1-32 threads in our later test, 1-4 threads yielded results with a MAPE of 0.505 while results from runs with 6-32 threads yielded results with a MAPE of 0.421. We are unaware of what may cause the different results.
